# Diet-induced obese mice are resistant to improvements in cardiac function resulting from short-term adropin treatment

**DOI:** 10.1101/2021.10.15.464576

**Authors:** Dharendra Thapa, Bingxian Xie, Bellina A.S. Mushala, Manling Zhang, Janet R. Manning, Paramesha Bugga, Michael W. Stoner, Michael J. Jurczak, Iain Scott

## Abstract

Previous studies have shown that treatment with recombinant adropin, a circulating peptide secreted by the liver and brain, restores glucose utilization in the hearts of diet-induced obese mice. This restoration of fuel substrate flexibility, which is lost in obese and diabetic animals, has the potential to improve contractile function in the diabetic heart. Using an *ex vivo* approach, we examined whether short-term adropin treatment could enhance cardiac function in a mouse model of diet-induced obesity. Our study showed that acute adropin treatment reduces inhibitory phosphorylation of pyruvate dehydrogenase in primary neonatal cardiomyocytes, and leads to moderate improvements in *ex vivo* cardiac function in mice fed a low fat diet. Conversely, short-term exposure to adropin led to a small decrease in cardiac function in mice fed a long-term high fat diet. Insulin treatment did not significantly alter cardiac function in adropin treated hearts from either low or high fat diet mice, however acute adropin treatment did moderately restore downstream insulin signaling in high fat diet fed mice. Overall, these data suggest that in an *ex vivo* setting, acute adropin treatment alone is not sufficient to promote improved cardiac function in obese animals.

## INTRODUCTION

Adropin is a short circulating peptide hormone produced in the liver and brain (Kumar et al., 2008). Once cleaved from its propeptide form, circulating adropin regulates metabolic function in a number of tissues, including the liver, brain, skeletal muscle, and the cardiovascular system (see, *e.g*., Kumar et al. 2008; Lovren et al., 2010; Gao et al., 2014; Gao et al., 2015; Stein et al., 2016; Thapa et al., 2019; Altamimi et al., 2019). Early studies demonstrated that long-term exposure to a high fat diet in mice resulted in decreased levels of circulating adropin (Kumar et al., 2008). Subsequent studies from the same group showed that restoration of adropin levels in diabetic mice, using either transgenic over-expression or treatment with recombinant peptide, led to a reversal in hyperglycemia, and the restoration of glucose oxidation in formerly insulin-resistant tissues such as skeletal muscle (Gao et al., 2014; Gao et al., 2015).

The ability of adropin to restore glucose oxidation in striated muscle from diabetic animals raised the possibility that it may have a beneficial effect on cardiac function in models of diabetic cardiomyopathy. While previous studies have shown that adropin can indeed improve glucose utilization in the hearts of both lean and diet-induced obese mice (Altamimi et al., 2019; Thapa et al., 2019), its effect on cardiac contractile function in obese animals remains unclear. In this study, we used an *ex vivo* isolated working heart approach to determine whether acute adropin treatment would improve cardiac functional parameters in the hearts of both lean and obese mice. Our studies demonstrate that while short-term adropin improves cardiac function in lean mice *ex vivo*, it has a small negative effect on cardiac contractile function in diet-induced obese mice. This lack of improvement may be the result of impaired insulin signaling in *ex vivo* hearts from obese mice, which is only moderately improved in adropin-treated animals fed a long-term high fat diet.

## METHODS

### Animal Husbandry and Use

Animals were housed in the University of Pittsburgh animal facility under standard conditions with *ad libitum* access to water and food on a constant 12h light/12h dark cycle. Male control and diet-induced obese C57BL/6J mice were obtained from The Jackson Laboratory after 22 weeks of either standard low fat diet (LFD; 70% carbohydrate, 20% protein, 10% fat; Research Diets D12450B), or a high fat diet (HFD; 20% carbohydrate, 20% protein, 60%fat; Research Diets D12492). Mice were maintained on this diet at the University of Pittsburgh for two weeks to acclimatize after transport prior to experimental use. At the end of 24 week LFD or HFD feeding regimens, mice received three I.P. injections of either vehicle (sterile PBS) or adropin (450 nmol/kg) over two days on a schedule described in Figure 2. After the second injection, mice were fasted overnight with free access to water. After the third injection, mice were euthanized and hearts rapidly excised for experimental use. Experiments were conducted in compliance with National Institutes of Health guidelines, and followed procedures approved by the University of Pittsburgh Institutional Animal Care and Use Committee.

### Neonatal Cardiomyocyte Isolation

Neonatal cardiomyocytes were isolated by collagenase disassociation from hearts obtained from P1-P3 rats. Cells were pre-plated to remove non-cardiomyocyte cells, and purified cardiomycoytes were seeded on collagen plates for 48 hours prior to experimental use. Cells were treated with vehicle (PBS) or adropin (0.5 μg/mL) for 24 hours, and then harvested for biochemical analysis.

### Protein Isolation and Western Blotting

Cardiac tissues were minced and lysed in CHAPS buffer (1% CHAPS, 150 mM NaCl, 10 mM HEPES, pH 7.4) on ice for ~2 hours. Homogenates were spun at 10,000 *g*, and supernatants collected for western blotting. Protein lysates were prepared in LDS sample buffer, separated using SDS/PAGE 4-12% or 12% Bis-Tris gels, and transferred to nitrocellulose membranes. Protein expression was analyzed using the following primary antibodies: mouse PDK4 (Abcam, catalog number ab110336, 1:1000), rabbit PDH (Cell Signaling, catalog number 2784, 1:1000), rabbit phospho-PDH Ser 293 (Cell Signaling, catalog number 31866, 1:1000), rabbit Tubulin (Cell Signaling, catalog number 2125, 1:5000), rabbit AKT (Cell Signaling, catalog number 9272, 1:1000), rabbit phospho-AKT Ser 473 (Cell Signaling, catalog number 4060, 1:1000), rabbit GSK-3β (Cell Signaling, catalog number 9315, 1:1000), rabbit GSK-3β Ser 9 (Cell Signaling, catalog number 5558, 1:1000). Fluorescent anti-mouse or anti-rabbit secondary antibodies (red, 700 nm; green, 800 nm) from Li-Cor were used to detect expression levels. Protein densitometry was measured using Image J software (National Institutes of Health, Bethesda, MD).

### Gene Expression Analysis

RNA was extracted from cells using RNEasy kit (Qiagen). cDNA was generated with 500 ng-1 μg of RNA using Maxima Reverse Transcriptase (ThermoFisher). Quantitative PCR (qPCR) was performed using SYBR-Green (ThermoFisher) reagent with primers for *Ppargc1a, Cd36, Cpt1b, Pdk4*, and *Gapdh* (Qiagen).

### Isolated Working Heart Analysis

Cardiac *ex vivo* function was calculated using a Harvard Apparatus ISHR isolated working heart system as previously described (Manning et al., 2019). Hearts from anesthetized mice were rapidly excised and cannulated via the aorta in warm oxygenated Krebs-Henseleit buffer (118 mM NaCl, 25 mM NaHCO_3_, 0.5 mM Na-EDTA [disodium salt dihydrate], 5 mM KCl, 1.2 mM KH_2_PO_4_, 1.2 mM MgSO_4_, 2.5 mM CaCl_2_, 11 mM glucose). Retrograde (i.e. Langendorff) perfusion was initiated to blanch the heart, maintained at a constant aortic pressure of 50 mmHg with a peristaltic pump through a Starling resistor. A small incision was next made in the pulmonary artery to allow perfusate to drain, and the heart was paced at a rate slightly higher than endogenous (~360-500 bpm). The left atrium was then cannulated via the pulmonary vein, and anterograde perfusion was initiated with a constant atrial pressure of 11 mmHg against an aortic workload of 50 mmHg. Left ventricle pressure was measured via Mikro-tip pressure catheter (Millar) carefully inserted into the LV through the aorta. The work-performing heart was permitted to equilibrate for 30 minutes to establish baseline functional parameters. After baseline measurements were completed, hearts were exposed to 0.1 U/L insulin for 10 minutes, after which hearts were either used for measurements of insulin-stimulated cardiac function, or snap-frozen for biochemical analyses. Mouse hearts that failed to equilibrate and/or function after cannulation were excluded from the working heart analysis.

### Statistical Analysis

Graphpad Prism software was used to perform statistical analyses. Means ± SEM were calculated for all data sets. Data were analyzed using either one-way or two-way ANOVA with Dunnett’s post-hoc multiple comparison testing to determine differences between treatment and feeding groups. Data were analyzed with two-tailed Student’s T-Tests to determine differences between single variable groups. *P* < 0.05 was considered statistically significant.

## RESULTS

### Adropin treatment reduces Pdk4 gene expression and inhibitory PDH phosphorylation in neonatal cardiomyocytes

We first examined the impact of adropin treatment on metabolic gene expression in primary neonatal cardiomyocytes. Exposure to adropin for 24 hours had no discernable effect on the expression of genes involved in fatty acid oxidation, including *Ppargc1a, Cd36*, and *Cpt1b* (Figure 1A-C). In contrast, *Pdk4* gene expression, a negative regulator of pyruvate dehydrogenase (PDH) activity, was significantly decreased by exposure to adropin (Figure 1D). The decrease in *Pdk4* gene expression resulting from adropin treatment was matched at the protein level (Figure 1E), which led to a significant decrease in inhibitory PDH phosphorylation at Ser 293. Based on these results, we conclude that adropin treatment is likely to improve glucose utilization in cardiomyocytes under normal nutrient conditions, in concordance with previous studies in H9c2 cells (Thapa et al., 2018) and mouse hearts (Altamimi et al., 2019).

**Figure 1.**
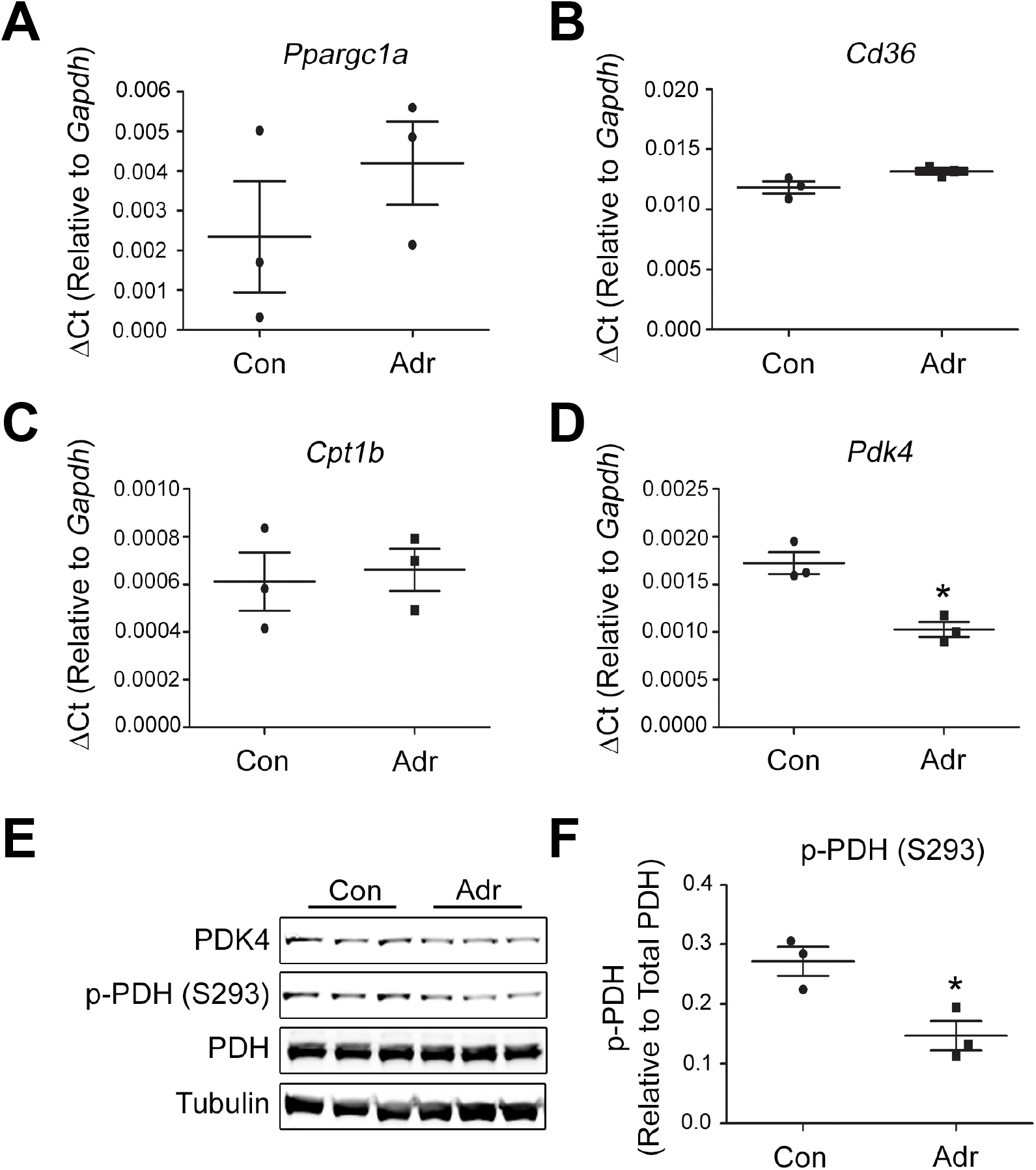
Adropin reduces PDK4 expression and inhibitory PDH phosphorylation in rat neonatal cardiomyocytes. (**A-D**) Adropin treatment significantly reduced Pdk4 gene expression in rat neonatal cardiomyocytes without affecting fatty acid oxidation pathway enzymes. (**E-F**) Adropin treatment reduced PDK4 protein expression, leading to significant reductions in inhibitory phosphorylation of pyruvate dehydrogenase (PDH). Con = control, Adr = Adropin. N = 3, * = *P* < 0.05 (Student’s T-Test).

### Exposure to a long-term high fat limits improvements in cardiac function driven by acute adropin treatment

We previously demonstrated that acute adropin treatment allows insulin-resistant, pre-diabetic mouse hearts to resume the use of glucose as a fuel substrate (Thapa et al., 2019). However, this study did not address whether improved glucose use led to improvements in cardiac contractility and workload. Therefore, we next examined whether a short-term adropin treatment regimen (Figure 2A) would result in increased cardiac function using an *ex vivo* isolated working heart approach. After 24 weeks of a high fat diet (HFD), there was an increase in body weight that was not affected by short-term adropin treatment (Figure 2B-C). At the end of the HFD exposure, there was a minor increase in contractility in vehicle-treated mice relative to their low fat diet (LFD) controls, along with a trend towards increased relaxation and cardiac output (Figure 3A-D). As shown previously by Altamimi et al. (2019), short-term adropin treatment led to an increase in all functional parameters (contractility, relaxation, workload, and output) in LFD mice (Figure 3A-D). In contrast, treatment of HFD mice with adropin led to a moderate decrease in cardiac function across the group relative to their vehicle-treated controls (Figure 3A-D). Based on these results, we conclude that acute adropin exposure in obese mice, in an *ex vivo* context, leads to an unexpected decrease in cardiac function.

**Figure 2.**
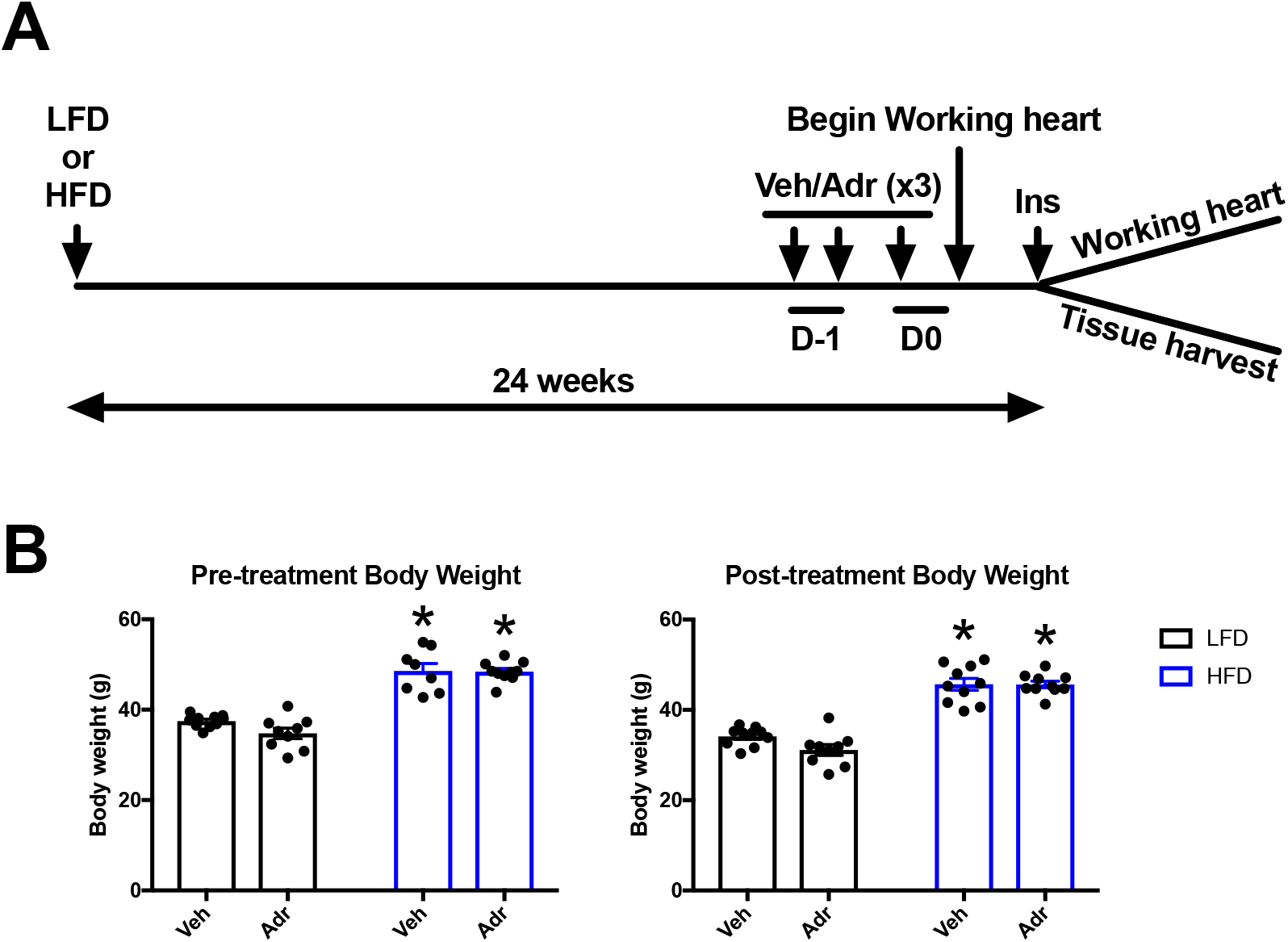
Schematic of *in vivo/ex vivo* experimental plan. (**A**) Male C57BL6/J mice aged six weeks were placed on a low fat diet (LFD; 10% fat) or high fat diet (HFD; 60% fat) for 24 weeks (N = 10 per group). On the day prior to organ harvest, mice received twice-daily IP injections of either vehicle (Veh; sterile PBS) or adropin (Adr; 450 nmol/kg in sterile PBS), and were then fasted overnight. On the morning of experiments, mice received a final IP injection of Veh or Adr, before hearts were rapidly excised and cannulated for isolated working heart measurements of cardiac function. After basal functional parameters were measured, hearts were infused with insulin, and randomly assigned to either further working heart analysis (N = 5 per group), or immediately snap-frozen for biochemical analysis (N = 5 per group). (B) Pre- and Post-treatment body weights of LFD and HFD mice. N = 10, * = *P* < 0.05 vs. LFD Veh group (Two-way ANOVA with Dunnett’s Post-Hoc Test).

**Figure 3.**
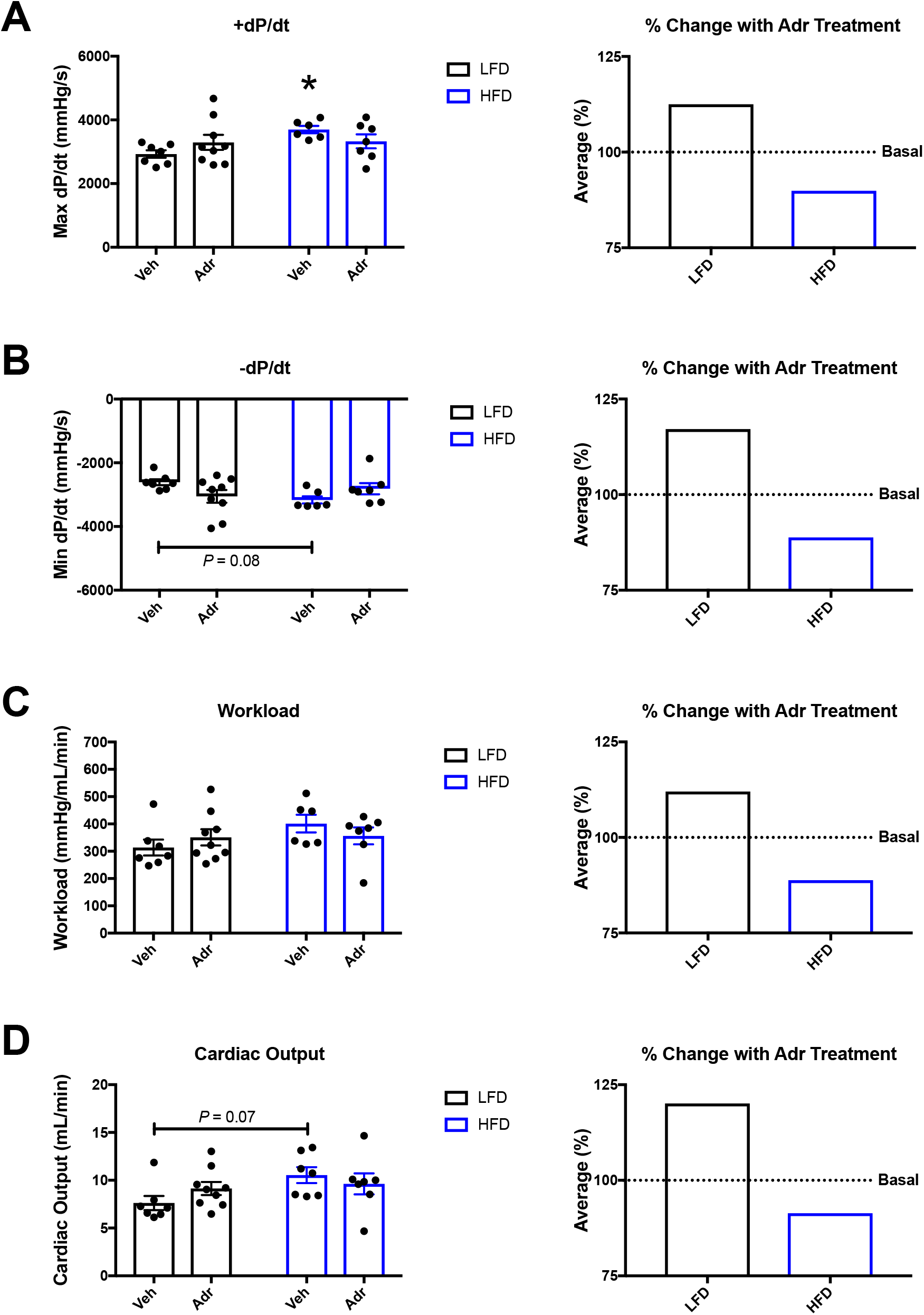
Exposure to a high fat diet inhibits adropin-driven improvements in cardiac function. (**A-D**) Vehicle treated (Veh) mice placed on a high fat diet (HFD) displayed a significant increase in contractility, and trends toward increased relaxation and cardiac output, relative to low fat diet (LFD) controls. Adropin (Adr) treatment in LFD mice led to moderate increases in all cardiac functional parameters, however this effect was reversed in mice exposed to a HFD. N = 6-9, * = *P* < 0.05 vs. LFD Veh group (Two-way ANOVA with Dunnett’s Post-Hoc Test).

### Insulin stimulation does not restore improvements in cardiac function after acute adropin treatment in high fat diet-exposed mice

Insulin stimulation leads to a shift towards glucose oxidation in adropin-treated lean mice (Altamimi et al., 2019) *ex vivo*, and in obese mice under hyperinsulinemic-euglycemic clamp conditions *in vivo* (Thapa et al., 2019). Therefore, we examined whether insulin stimulation would reverse the loss of cardiac function in adropin-treated mice using our isolated working heart approach. In LFD mice, insulin stimulation of adropin treated animals again resulted in a moderate average increase in cardiac functional parameters (Figure 4A-D). However, the ability of insulin to drive glucose oxidation in the hearts of HFD adropin-treated mice did not result in improved cardiac contractility or output (Figure 4A-D). Based on these results, we conclude that insulin stimulation is not sufficient to restore contractility in adropin-treated HFD mouse hearts *ex vivo*.

**Figure 4.**
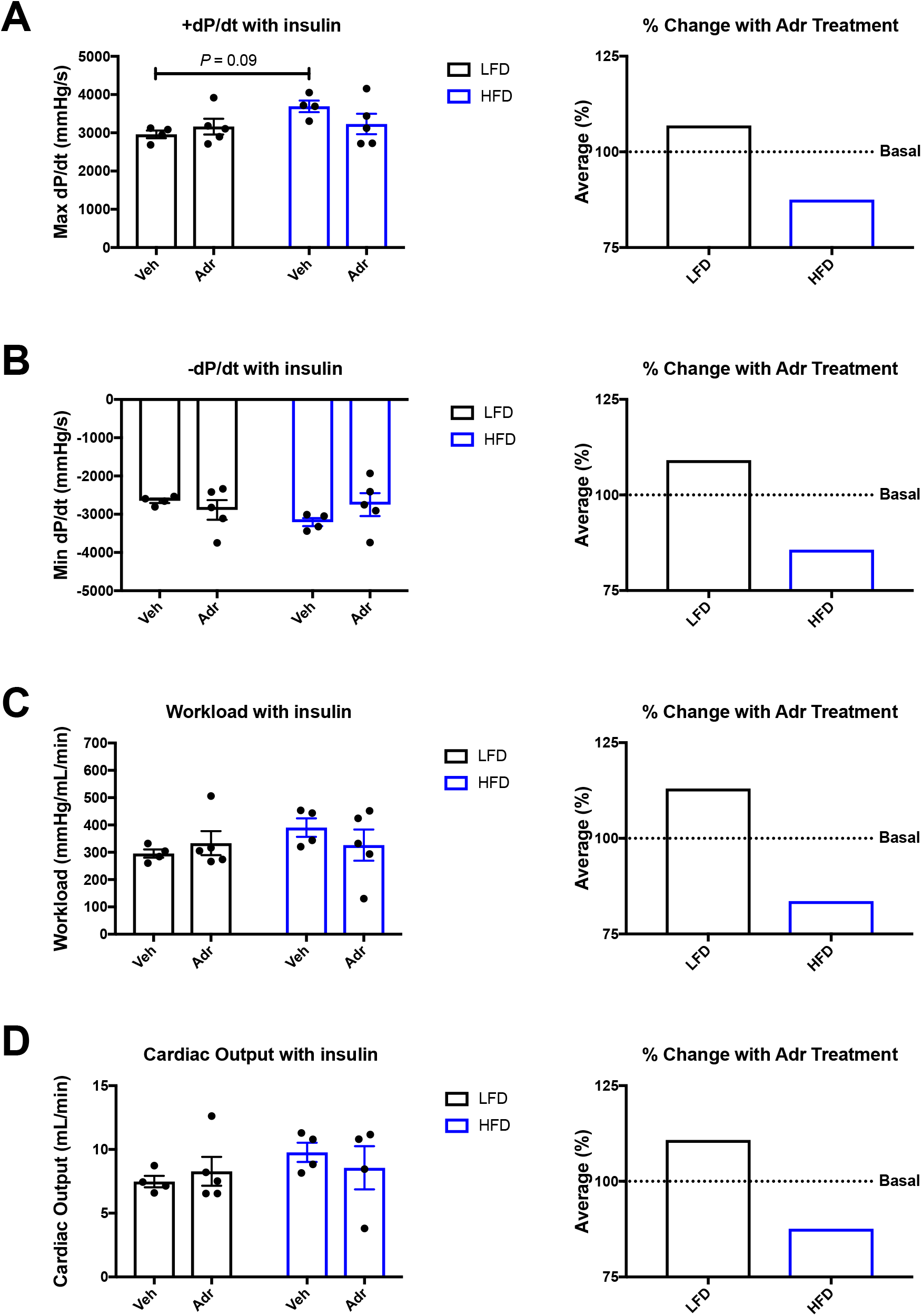
Insulin exposure does not reverse the loss of adropin-driven improvements in cardiac function in mice exposed to a high fat diet. (**A-D**) As in the unstimulated state (Figure 3), mice on a low fat diet (LFD) displayed moderate increases in cardiac functional outcomes after insulin stimulation following acute adropin (Adr) treatment. Conversely, mice exposed to a high fat diet (HFD) displayed decreased cardiac function in response to insulin stimulation after acute adropin treatment. N = 4-5, * = *P* < 0.05 vs. LFD Vehicle (Veh) group (Two-way ANOVA with Dunnett’s Post-Hoc Test).

### Acute adropin treatment moderately improves cardiac insulin signaling in high fat diet-exposed mice

In lean mice, adropin treatment leads to an increase in cardiac insulin sensitivity, as measured by the induction of cellular signaling pathways (Altamimi et al., 2019). To understand if exposure to a HFD was blocking the ability of adropin to induce insulin signaling pathways, we first examined AKT activation in hearts from lean and obese mice after insulin exposure. In vehicle-treated LFD mice, insulin stimulation led to a significant increase in AKT activation, as measured by phosphorylation at Ser 473 (Figure 5A). However, in both vehicle- and adropin-treated HFD mice, there was no significant induction of AKT signaling after insulin exposure (Figure 5A). We next examined insulin signaling downstream of AKT, by measuring phosphorylation of GSK-3β at Ser 9. As with AKT, insulin exposure led to a significant induction of GSK-3β phosphorylation in vehicle-treated LFD mice (Figure 5B). While vehicle-treated HFD mice showed no response to insulin (in keeping with AKT, above), adropin-treated HFD mice displayed a significant increase in GSK-3β phosphorylation at Ser 9 (Figure 5B). Based on these results, we conclude that acute adropin treatment has a moderate positive effect on downstream insulin signaling in obese mice.

**Figure 5.**
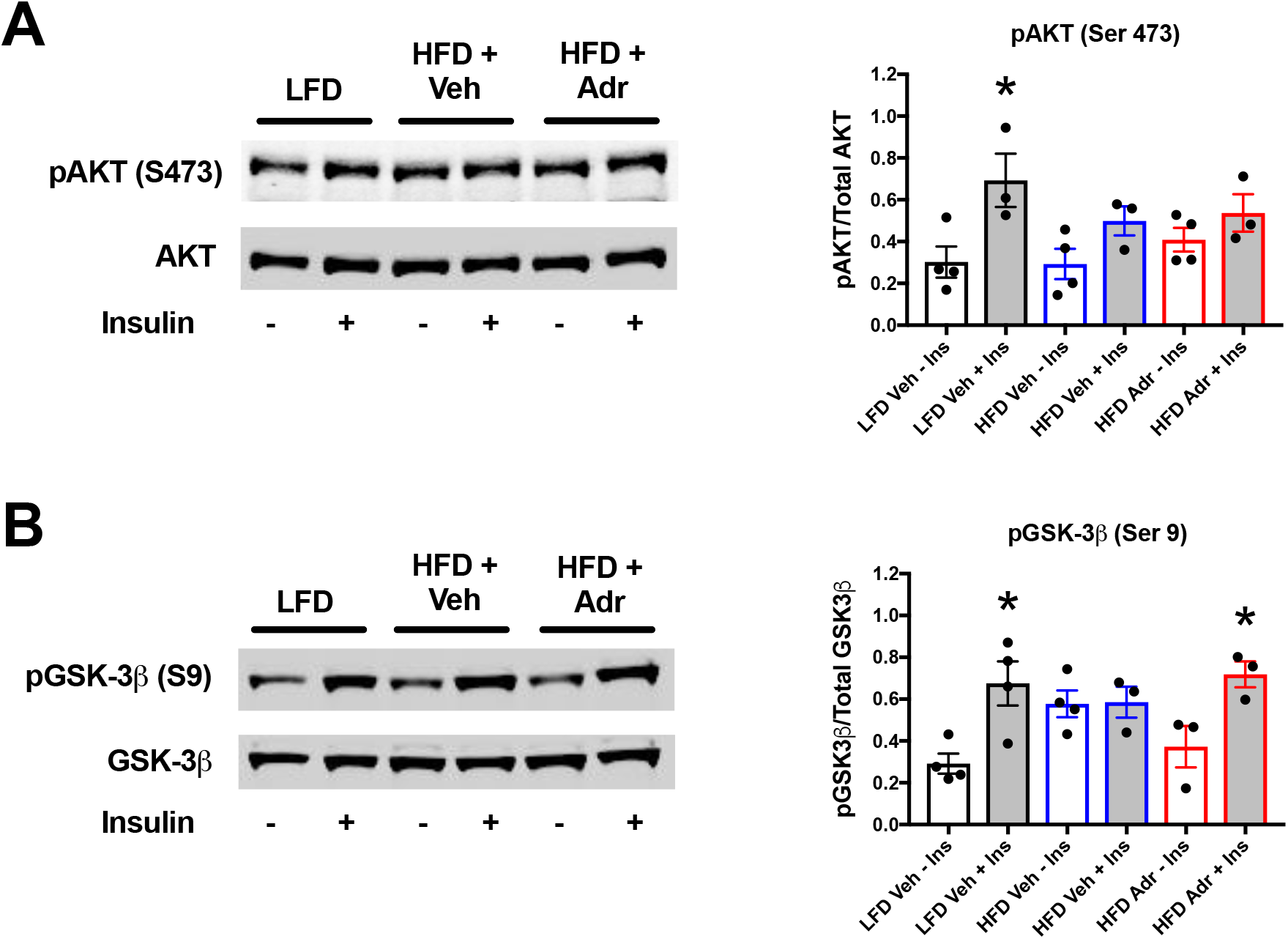
Acute adropin treatment moderately improves downstream insulin signaling in mouse hearts exposed to a high fat diet. (**A-B**) Mice on a low fat diet (LFD) were responsive to insulin stimulation, as shown by phosphorylation of AKT at Ser 473. Conversely, both vehicle (Veh) and adropin (Adr) treated hearts did not shown significant changes in AKT phosphorylation after exposure to a high fat diet (HFD). The same pattern was observed in downstream AKT signaling (phosphorylation of GSK-3β at Ser 9) in vehicle-treated HFD mice after insulin stimulation. However, adropin treatment restored insulin-mediated GSK-3β phosphorylation in HFD mice under the same conditions. N = 3-4, * = *P* < 0.05 vs. LFD Veh group (One-way ANOVA with Dunnett’s Post-Hoc Test).

**Figure 6.**
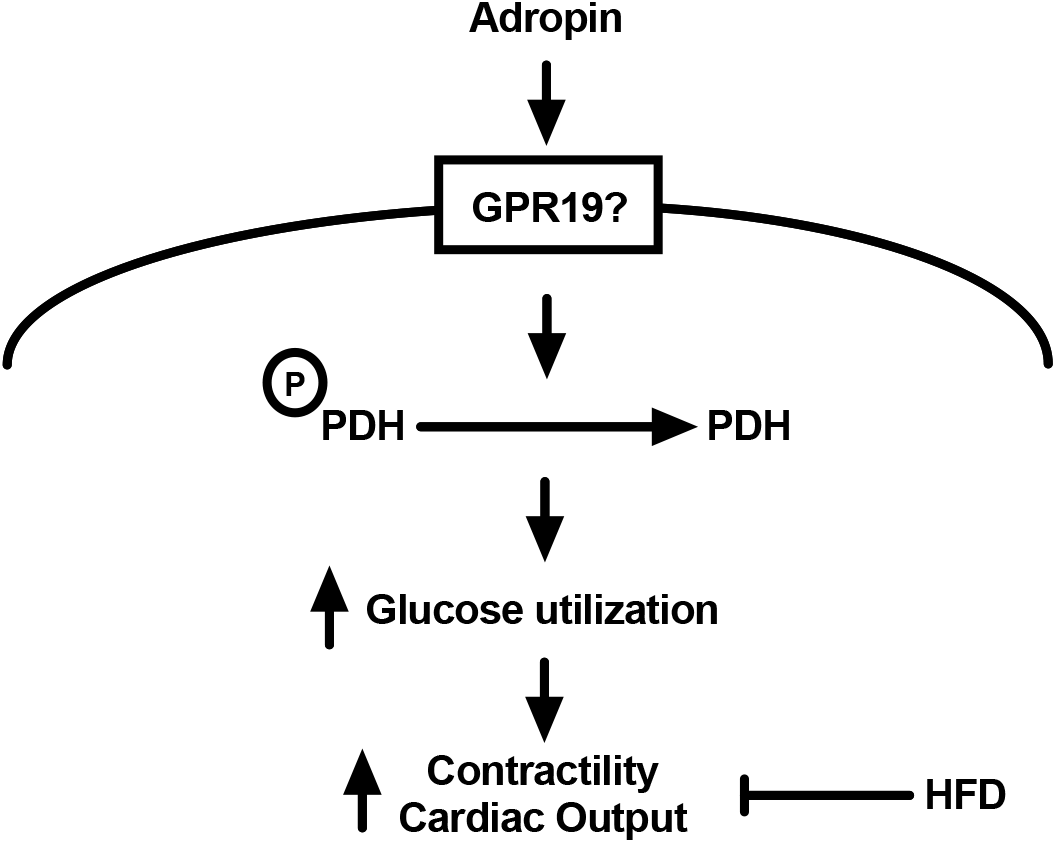
Model of acute adropin treatment on cardiac function in mice exposed to a high fat diet. Short-term adropin exposure, potentially signaling through the cell surface receptor GPR19, results in decreased inhibitory pyruvate dehydrogenase (PDH) phosphorylation and increased cardiomyocyte glucose utilization. In animals on a low fat diet, this leads to increased cardiac contractility and output, however the effect is lost after exposure to a long-term high fat diet (HFD).

## DISCUSSION

In keeping with previous studies, we show that short-term adropin treatment reduces inhibitory phosphorylation of PDH in primary cardiomyocytes *in vitro*, and increases overall cardiac function in the hearts from lean mice *ex vivo*. For the first time, we show that acute adropin treatment has a small detrimental effect on cardiac function in obese mouse hearts, when examined in an *ex vivo* context. Furthermore, we show that while adropin treatment can moderately improve downstream insulin signaling in mice fed a high fat diet, it does not restore proximal insulin signaling through AKT.

Studies on the role of adropin in cardiac energy metabolism were prompted by reports of its function in skeletal muscle. Elegant work by the Butler group demonstrated that acute adropin treatment downregulated genes involved in fatty acid oxidation, reduced inhibitory PDH phosphorylation via reductions in *Pdk4* expression, and restored insulin signaling in the skeletal muscle of diet-induced obese mice (Gao et al., 2014; Gao et al., 2015). Importantly, in addition to showing that acute adropin treatment reduced whole-body hyperglycemia, they used indirect calorimetry to show that adropin shifted oxidation preferences from fat to glucose in obese mice (Gao et al., 2015). Follow-up studies in the heart demonstrated that all of these same metabolic pathways were operable, and that acute adropin treatment could promote glucose utilization in the hearts of both lean and obese mice (Altamimi et al., 2019; Thapa et al., 2019).

While cardiac function was improved by acute adropin treatment in lean mice *ex vivo* (Altamimi et al., 2019), its effect on the obese mouse heart is less clear. Thapa et al. (2019) showed that three days of adropin treatment had little effect on systolic function, with a non-significant trend towards improved diastolic function in adropin-treated obese mice *in vivo*. Our findings in this study suggest that acute adropin exposure in mice exposed to a long-term HFD is moderately detrimental in terms of cardiac function when measured *ex vivo* (Figures 3,4). The mechanism underlying this decrease in function is unclear, but may be the result of several factors. Firstly, this short-term adropin treatment did not fully re-establish insulin signaling in the HFD mouse hearts (Figure 5), which may have left these hearts mildly energy starved relative to their LFD controls. However, while the perfusate contained only glucose, isolated working hearts from rodents can maintain fatty acid oxidation from endogenous triglyceride pools for at least 60 minutes (Saddik and Lopaschuk, 1991), suggesting that energy supply *per se* may not be the main cause of functional decline. Secondly, the adropin treatment regimen used here (three injections over two days) was shorter than our previous *in vivo* studies (five injections over three days; Thapa et al., 2019), and this may have abrogated its biological response. Thirdly, the isolated working heart model used necessarily operates in the absence of neurohormonal stimulation (reviewed in Ghionzoli et al. 2021), and our understanding of the interplay between adropin and other hormonal regulators of cardiac function remains incomplete. Finally, previous studies in non-diabetic failing hearts have shown that blocking cardiac glucose uptake via insulin resistance may be cardioprotective, by preventing glucotoxicity from incomplete glucose metabolism (Taegtmeyer et al., 2013). While the short-term switch to increased glucose use driven by adropin treatment may potentiate such a response, this may be viewed as a less likely outcome, as glucose oxidation appears to be complete in both lean and obese mice after adropin treatment (Altamimi et al., 2019; Thapa et al., 2019).

To address these questions, future work will need to focus on two main factors. Firstly, given the uncertainty surrounding potential lack of neurohormonal stimulation, further *in vivo* studies of cardiac fuel metabolism and function will need to be performed in both lean and obese mice after adropin treatment. Secondly, given that structural changes (dilation, hypertrophy, etc.) may occur after extended periods of high fat feeding in mice, it is unlikely that a short-term adropin treatment regimen will be sufficient to reverse these outcomes. As such, future studies will need to examine whether long-term adropin replacement is required to improve cardiac functional outcomes in obese and diabetic animals.

## FUNDING

This work was supported by: NIH/NHLBI K99/R00 award (HL146905) to DT; NIGMS T32 award (GM133332) to B.A.S.M; and NIH/NHLBI R01 awards (HL132917 & HL147861) to IS. The University of Pittsburgh Center for Metabolism and Mitochondrial Medicine (C3M) is supported by a Pittsburgh Foundation award (MR2020 109502) to MJJ.

## AUTHOR CONTRIBUTIONS

DT, BX, MJJ, and IS designed the experiments. DT, BX, MZ, MWS, and JRM performed the experiments. DT, BX, and IS analyzed the data. IS produced the figures. PB and BASM provided critical input into the manuscript and discussion. DT and IS wrote and edited the manuscript.

## DISCLOSURES

None.

## Notes

### Competing Interest Statement

The authors have declared no competing interest.

## REFERENCES

Altamimi TR, Gao S, Karwi QG, Fukushima A, Rawat S, Wagg CS, Zhang L, Lopaschuk GD. Adropin regulates cardiac energy metabolism and improves cardiac function and efficiency. Metabolism. 2019 Sep;98:37–48. doi: 10.1016/j.metabol.2019.06.005. Epub 2019 Jun 14. PMID: 31202835.

Gao S, McMillan RP, Jacas J, Zhu Q, Li X, Kumar GK, Casals N, Hegardt FG, Robbins PD, Lopaschuk GD, Hulver MW, Butler AA. Regulation of substrate oxidation preferences in muscle by the peptide hormone adropin. Diabetes. 2014 Oct;63(10):3242–52. doi: 10.2337/db14-0388. Epub 2014 May 21. PMID: 24848071; PMCID: PMC4171656.

Gao S, McMillan RP, Zhu Q, Lopaschuk GD, Hulver MW, Butler AA. Therapeutic effects of adropin on glucose tolerance and substrate utilization in diet-induced obese mice with insulin resistance. Mol Metab. 2015 Jan 17;4(4):310–24. doi: 10.1016/j.molmet.2015.01.005. PMID: 25830094; PMCID: PMC4354928.

Ghionzoli N, Gentile F, Del Franco AM, Castiglione V, Aimo A, Giannoni A, Burchielli S, Cameli M, Emdin M, Vergaro G. Current and emerging drug targets in heart failure treatment. Heart Fail Rev. 2021 Jul 17. doi: 10.1007/s10741-021-10137-2. Epub ahead of print. PMID: 34273070.

Kumar KG, Trevaskis JL, Lam DD, Sutton GM, Koza RA, Chouljenko VN, Kousoulas KG, Rogers PM, Kesterson RA, Thearle M, Ferrante AW Jr, Mynatt RL, Burris TP, Dong JZ, Halem HA, Culler MD, Heisler LK, Stephens JM, Butler AA. Identification of adropin as a secreted factor linking dietary macronutrient intake with energy homeostasis and lipid metabolism. Cell Metab. 2008 Dec;8(6):468–81. doi: 10.1016/j.cmet.2008.10.011. PMID: 19041763; PMCID: PMC2746325.

Lovren F, Pan Y, Quan A, Singh KK, Shukla PC, Gupta M, Al-Omran M, Teoh H, Verma S. Adropin is a novel regulator of endothelial function. Circulation. 2010 Sep 14;122(11 Suppl):S185–92. doi: 10.1161/CIRCULATIONAHA.109.931782. PMID: 20837912.

Manning JR, Thapa D, Zhang M, Stoner MW, Traba J, McTiernan CF, Corey C, Shiva S, Sack MN, Scott I. Cardiac-specific deletion of GCN5L1 restricts recovery from ischemia-reperfusion injury. J Mol Cell Cardiol. 2019 Apr;129:69–78. doi: 10.1016/j.yjmcc.2019.02.009. Epub 2019 Feb 15. PMID: 30776374; PMCID: PMC6486843.

Saddik M, Lopaschuk GD. Myocardial triglyceride turnover and contribution to energy substrate utilization in isolated working rat hearts. J Biol Chem. 1991 May 5;266(13):8162–70. PMID: 1902472.

Stein LM, Yosten GL, Samson WK. Adropin acts in brain to inhibit water drinking: potential interaction with the orphan G protein-coupled receptor, GPR19. Am J Physiol Regul Integr Comp Physiol. 2016 Mar 15;310(6):R476–80. doi: 10.1152/ajpregu.00511.2015. Epub 2016 Jan 6. PMID: 26739651; PMCID: PMC4867374.

Taegtmeyer H, Beauloye C, Harmancey R, Hue L. Insulin resistance protects the heart from fuel overload in dysregulated metabolic states. Am J Physiol Heart Circ Physiol. 2013 Dec;305(12):H1693–7. doi: 10.1152/ajpheart.00854.2012. Epub 2013 Oct 4. PMID: 24097426; PMCID: PMC3882545.

Thapa D, Stoner MW, Zhang M, Xie B, Manning JR, Guimaraes D, Shiva S, Jurczak MJ, Scott I. Adropin regulates pyruvate dehydrogenase in cardiac cells via a novel GPCR-MAPK-PDK4 signaling pathway. Redox Biol. 2018 Sep;18:25–32. doi: 10.1016/j.redox.2018.06.003. Epub 2018 Jun 9. PMID: 29909017; PMCID: PMC6008287.

Thapa D, Xie B, Zhang M, Stoner MW, Manning JR, Huckestein BR, Edmunds LR, Mullett SJ, McTiernan CF, Wendell SG, Jurczak MJ, Scott I. Adropin treatment restores cardiac glucose oxidation in pre-diabetic obese mice. J Mol Cell Cardiol. 2019 Apr;129:174–178. doi: 10.1016/j.yjmcc.2019.02.012. Epub 2019 Feb 26. PMID: 30822408; PMCID: PMC6486841.

